# M^6^A reader protein YTHDF3 regulates cardiomyocyte death and atrophy by modulating the alternative splicing program

**DOI:** 10.1101/2025.04.15.648887

**Authors:** Aakash Gaur, Sakshi Chaudhary, Rakesh Kumar Sharma, Samprikta Kundu, E Ganesh, Renu Kumari, Sandhya Singh, Shakti Prakash, Kalyan Mitra, Shailesh Kumar, Shashi Kumar Gupta

## Abstract

**Background:** The functional impact of m^6^A modifications on RNA is governed by reader proteins that read the m^6^A marks and process the RNA accordingly. Recent studies have highlighted the importance of m^6^A reader proteins in the heart. However, the function of a reader protein *Ythdf3* in the heart remains completely unknown.

**Objective:** We aim to uncover the function of *Ythdf3* in the hearts.

**Methods& Results:** Here, we show that *Ythdf3* is indispensable for the heart, and its knockdown leads to cardiac cachexia evident by cardiomyocyte atrophy, death, and less dense myofibrils visualized by electron microscopy. We also found downregulation of *Ythdf3* in response to doxorubicin, and its overexpression rescues doxorubicin-induced cardiomyocyte atrophy and death. Contrary to primarily cytoplasmic localization, we found that *Ythdf3* localizes in the cardiomyocyte nucleus. Furthermore, using co-immunoprecipitation coupled with LC-MS/MS, we show that *Ythdf3* interacts with splicing proteins like DDX5 and HNRNPU and regulates alternative splicing in the heart. Mechanistically, we found that *Ythdf3* regulates the splicing of CaMKIIδ, and its knockdown leads to an increase in CaMKIIδA and CaMKIIδC isoforms while a decrease in CaMKIIδ9 isoform.

**Conclusion:** Collectively, *Ythdf3* knockdown in the adult heart leads to cardiac cachexia due to the alteration of the splicing program.

## Introduction

Chemical modification of RNA molecules is an essential post-transcriptional process to regulate gene expression and function [10]. Modifications found on RNA include N^6^-methyladenosine (m^6^A), pseudouridine (Ψ), N^5^-methylcytidine (m^5^C), N^1^-methyladenosine (m^1^A), N^4^-acetylcytidine (ac^4^C), dihydrouridine (D), and 2’-O-methyl ribose (Nm) [10]. Among the diverse modifications found on RNAs, m^6^A is the most prevalent RNA modification. M^6^A is found in all RNAs, including mRNAs, tRNAs, rRNAs, miRNAs, lncRNAs, and circular RNAs [14]. M^6^A modifications regulate RNA secondary structure, stability, translation, splicing, and localization [14].

M^6^A methylation is a reversible modification, added by methyltransferases or writers, mainly METTL3/14/16, and removed by demethylases or erasers, *FTO* and *ALKBH5* [14]. Cardiomyocyte-specific deletion of METTL3 in mice led to cardiac malfunction with aging and in response to pressure overload [7]. Cardiomyocyte-specific overexpression of METTL3 led to physiological hypertrophy and maintained cardiac function in response to pressure overload [7]. Another writer protein, METTL14, is downregulated in DCM (dilated cardiomyopathy), and its overexpression improves cardiac function [18]. METTL14 regulates pyroptosis in cardiomyocytes by regulating the expression of lncRNA TINCR in a m^6^A-dependent manner [18]. RNA m^6^A methylation is a reversible process, and demethylases can erase m^6^A methylation marks; therefore, demethylases are essential in controlling the temporal and spatial m^6^A methylation levels. A recent study showed concomitantly higher levels of m^6^A methylation and lower levels of demethylase FTO in the failing hearts of humans, pigs, and mice. AAV9-mediated overexpression of FTO improved cardiac function in a mouse model of myocardial infarction by demethylating contractile protein mRNAs [17]. Another demethylase, ALKBH5, has been shown to play an essential role in doxorubicin-induced cardiotoxicity [4]. Overexpression of ALKBH5 exacerbated doxorubicin-induced cardiotoxicity, while knockdown rescued the doxorubicin effects evident by changes in cardiac function, cardiac atrophy, and apoptosis [4]. These results highlight the critical function of m^6^A in maintaining cardiac homeostasis and survival. However, the functional impact of m^6^A modifications on RNA is driven by reader proteins that read the m^6^A marks and process the RNA accordingly.

YTH domain-containing proteins are one of the essential reader proteins recognizing m^6^A modifications on RNA and regulating their fate [20]. There are two sub-families of YTH proteins, YTHDC (YTHDC1 & YTHDC2) and YTHDF (YTHDF1, YTHDF2, and YTHDF3) [20]. Cardiac-specific knockdown of YTHDC1 in adult mice led to dilated cardiomyopathy due to dysregulation of Titin splicing [9]. Few recent studies have shown the translational regulatory role of YTHDF proteins in the heart. Recently, cardiomyocyte-specific-YTHDF1 knockout has been shown to develop hypertrophy and heart failure [11]. Loss of YTHDF1 led to a decline in the translation of several genes involved in membrane raft organization. On contrary to YTHDF1 translational promoting function, another YTHDF protein, YTHDF2, has been found to exert translation inhibiting function. Cardiomyocyte-specific deletion of YTHDF2 leads to cardiac hypertrophy and deteriorated cardiac function [12, 15]. Cardiac hypertrophy observed upon loss of YTHDF2 was due to increased translation of eukaryotic elongation factor EEF2 and MYZAP [12, 15]. Thus, cytoplasm-localized YTHDF proteins with their translational regulatory function regulate the cardiac proteome and aid in maintaining cardiac homeostasis. However, another member of YTHDF family, YTHDF3 has been relatively less studied in the heart with its function in adult heart remaining completely unknown.

Here, we have knockdown YTHDF3 in cardiomyocytes using shRNA delivered via AAV (Adeno-associated virus). Knockdown of YTHDF3 led to cardiomyocyte death and atrophy *in vitro* and *in vivo*. In contrast to its cytoplasmic localization in other cell types, YTHDF3 was located in the nucleus of cardiomyocytes. Using immunoprecipitation, the YTHDF3 interactome reveals enrichment for splicing regulators. We have found that YTHDF3 regulates alternative splicing of CaMKIIδ.

## Materials and methods

### Cell culture and Lentivirus production

H9c2 rat myoblast and Human embryonic kidney (HEK293T) cells were cultured in DMEM high glucose medium (Thermo Scientific, USA) with 10% FBS. For inhibition experiments in H9c2 cells, scramble and *Ythdf3* shRNA oligos were synthesized and cloned in pLKO-shRNA-eGFP-WPRE plasmid with AgeI and EcoRI (Thermo Scientific, USA) restriction enzymes. For lentivirus production, HEK293T cells were transfected using PEIMAX 40000 (Polyscience #49553-93-7) with pLKO shRNA and lentiviral helper plasmids (psPAX2 and pMD2.G). After the twenty-four-hour transfection, the medium was replaced with fresh 2.5% FBS containing DMEM medium, and lentiviral containing supernatant was collected at forty-eight, seventy-two, and ninety-six hours post medium change. The collected supernatant was precipitated with polyethylene glycol 8000(Sigma #P2139) and centrifuged at 2500*g. Concentrated lentiviral particles were dissolved in 1X phosphate buffer saline, pH 7.4. All the shRNA sequences are given in **Supplementary Table 1**.

### Adeno-associated viruses

*Ythdf3* and Scramble shRNAs were cloned in pAAV-U6 plasmids between AgeI and EcoRI (Thermo Scientific,USA) restriction enzyme sites for inhibition study in isolated cardiomyocytes and mouse hearts. The *Ythdf3* gene was amplified from mouse heart cDNA and cloned in pAAV-cTNT plasmid with EcoRI and BamHI restriction enzymes for overexpression. AAV was produced by an already-established protocol [16, 21]. Concisely, HEK293T cells were transfected using PEIMAX 40000 (Polyscience #49553-93-7) with AAV-U6 shRNAs or AAV-cTNT-*Ythdf3* plasmid with AAV helper plasmids. For serotypes 6 and 9, pDG6 and pDG9 helper plasmids were used, respectively. After twenty-four hours of transfection, the medium was changed with AAV production medium having high glucose DMEM, 1% FBS, 1%penicillin/streptomycin, 10mM HEPES (Sigma Aldrich, USA), 0.075% sodium bicarbonate and 1X Glutamax (Gibco, USA). Secreted AAV particles from the medium were precipitated using polyethylene glycol 8000 (SIGMA #P2139). Cells containing AAVs were processed by acidic lysis, and AAV particles were further precipitated with polyethylene glycol, collected after centrifugation at 2500*g, and dissolved in 1X phosphate buffer saline. PBS-dissolved AAV particles were also cleaned with chloroform for *in vitro* application. For *in vivo* application, chloroform purified AAV9 was ultra-purified using Optiprep (Sigma Aldrich, USA) density gradient ultracentrifugation (Beckman Coulter, USA). Ultra-purified AAV from 40% iodixanol fraction was collected by puncturing the ultracentrifugation tube. AAV particles were then concentrated and washed using Amicon 100 K (100 kDa) cut-off columns (Sigma Aldrich, USA). AAV9 and AAV6 were titrated by real-time PCR using U6 and cTNT promoter-based primers.

### Neonatal rat/mouse cardiomyocyte isolation

One to three days old, rat or mouse pups were sacrificed, and their hearts were explanted and chopped by scalpel blade. Chopped heart tissues were digested using Collagenase Type II at 1 mg/ml (Thermo Scientific, USA). After successful digestion, the enzymatic reaction was stopped using 20% FBS containing DMEM medium, and cells were passed through a 70 μm pore size cell strainer to remove undigested tissue. Cells were centrifuged at low speed (40*g) to get enriched cardiomyocyte fraction and pre-plated in 20% FBS-containing medium for ninety minutes. Nonadherent cells (cardiomyocytes) were collected and counted from the pre-plated plate using a hemocytometer.

For cell size and TUNEL experiments, eighty thousand and forty thousand cells were seeded in twenty-four and forty-eight well plates, respectively. After forty-eight hours’ cells were washed with 1X PBS, and 1% FBS containing DMEM was added. Cardiomyocytes were transduced with AAV6 at MOI 3*10^5^ -1*10^5^ for 5-7 days for different experimental setups. At the end of the experiment, cells were washed with 1XPBS three times and fixed with 4% paraformaldehyde solution for cell size and TUNEL experiments.

### Adult mouse cardiomyocyte isolation

A previously established protocol was used for adult cardiomyocyte isolation [1]. Concisely, the mouse was deeply anesthetized by isoflurane in a big petri dish. The mouse chest cavity was opened, and all the blood was removed from the body after cutting the descending aorta and inferior vena cava. The cardiac contraction was seized by slowly injecting 7ml of EDTA buffer from the right ventricle. Next, the heart was explanted by cutting outside the clamped ascending aorta using Reynolds forceps and transferred to a 60 mm dish containing EDTA buffer. Again, 5ml of EDTA buffer was injected from the apex of the left ventricle at a very slow speed(1ml/2-3min) to completely stop the contraction of the heart. The non-contractile heart was transferred to the next dish containing perfusion buffer, and 3-5ml of perfusion buffer was injected from the left ventricle apex to remove the remaining blood. Finally, the heart is transferred in a digestion buffer dish kept at a 37°C hot plate and digested multiple times with 30ml of digestion buffer per animal. After successful digestion of heart tissue, 5% FBS containing DMEM was added to stop the enzymatic reaction, and cells were passed through a cell strainer (100 μm pore size). Enriched cardiomyocyte fraction is obtained by low-speed centrifugation 90*g, and cells were seeded in laminin-coated 96 well plates and kept in a culture medium containing calcium for 5 hours. Adhered cells were carefully washed with 1X PBS and fixed with 0.1% paraformaldehyde for further experimental purposes.

### RNA and PCR

Total RNA was isolated from H9c2, neonatal rat cardiomyocytes, whole mouse heart, and isolated adult mouse cardiomyocytes using RNA Iso (Takara Bio, Japan). For cDNA synthesis, 1–2 μg of total RNA was reverse transcribed with random hexamers using PrimeScript 1st strand cDNA synthesis kit (Takara Bio, Japan) on Veriti 96-well thermocycler (Thermo Scientific, USA). Real-time amplification was performed using TB green Premix EXtaq (Takara Bio) as per the manufacturer’s protocol using target gene-specific forward and reverse primers (Sigma Aldrich and Barcode Biosciences) on QuantStudio 12 K Flex (Thermo Scientific, USA). For CaMKIIδ alternative splicing studies, primers were designed at two constitutive exons to amplify all the transcripts, and the amplified product was run on 6% DNA Polyacrylamide gel (DNA PAGE). After the run, the gel was stained with Ethidium bromide and visualized in the gel doc system (Bio-Rad Gel Doc XR+). All the primer sequences are given in **Supplementary Table 2**.

### Western blotting

Cardiomyocyte pellets were lysed in cell lysis buffer (1X RIPA) on ice and sonicated, while mouse heart tissues were crushed in liquid nitrogen, dissolved in 1X RIPA buffer, and centrifuged at 12000*g. Protein concentration was estimated using the BCA kit (Thermo Scientific, USA). 20-60μg protein was loaded on 10% SDS Polyacrylamide gel, and the resolved protein was transferred to PVDF membrane (Bio-Rad, USA) using a Mini PROTEAN Tetra cell (Bio-Rad, USA) at 4°C temperature for 2 hours. PVDF membrane was then blocked by 5% bovine serum albumin and incubated with primary antibody of YTHDF3 (Invitrogen, #PA597034.), YTHDF3 (Abclonal, #8395), Ddx5 (Abclonal, #A11339), HNRNPU (Abclonal, #A4257), GAPDH (#MA515738 Thermo Scientific, USA), Cleaved Caspase-3 (# Sigma-Aldrich AB3623), and ACTB (#A00730-100 Genscript, USA) for overnight at 4°C. The next day, the membrane was washed three times with TBST buffer and incubated with secondary antibodies linked to HRP (Thermo Scientific, USA). For protein size, a reference ladder was used (Gene to Protein, India, and Bio-Rad).

### Cell size measurements

Paraformaldehyde fixed neonatal rat cardiomyocytes were permeabilized with 0.1% Triton-X-100 and stained with mouse α-sarcomeric actinin (#MA1-22,863 Thermo Scientific, USA) in a 1:1000 ratio overnight at 4°C temperature. The next day, cells were incubated with Alexa flour labeled anti-mouse secondary antibody. DAPI was used to stain the nucleus. Cardiomyocyte images were captured at 20L×Lobjective using Leica DMI 6000 B (Leica Microsystems, USA). ImageJ software was used to measure the cardiomyocyte area. All cell size experiments were repeated three times (biological replicate) with 2-3 wells each. More than 100 cells were counted from each well. Isolated adult cardiomyocytes from mouse hearts were plated in a 10mm culture dish, and images were captured using an inverted light microscope (Nikon, USA). Myocyte area, length, and width were measured using ImageJ software.

Cryo-preserved heart sections were cut using Cryotome (Thermo Scientific, USA). Cryosections were then fixed and permeabilized. Sections were further stained with wheat germ agglutinin (#MP00831 Thermo Scientific, USA) and DAPI. All images were taken at 20L×Lobjective using Leica DMI 6000 B (Leica Microsystems, USA). Cell cross-sectional area was measured using ImageJ software.

### Subcellular fractionation

Nuclear and cytoplasmic protein fractions of H9c2 cells were isolated using already published methods [5]. In brief, H9c2 cells were Trypsinized and pelleted down after centrifugation at 1000 RPM. The cell pellet was lysed with 400ul of NP-40 lysis buffer for 5 min on ice, gently loaded on 24% sucrose buffer in 2ml centrifuge tube, and centrifuged at 3500*g for 10 min. The upper layer, which had cytoplasmic content, was separated, and the nuclear pellet was washed and collected for western blotting and pull-down experiments.

### TUNEL (terminal deoxynucleotidyl transferase dUTP nick end labeling) staining

For TUNEL staining, cardiomyocytes were fixed with 4% PFA prepared in nuclease-free water and permeabilized on ice with 0.1% Triton-x-100. Labeling solution with terminal deoxynucleotidyl transferase (TdT) was mixed as per the manufacturer’s In Situ Cell Death Detection Kit (Roche) protocol, and cells were incubated for 45 min at 37°C temperature. Next, the reaction mixture was removed, and cells were stained with α-sarcomeric actinin and DAPI. Images were captured at 20L×Lobjective using Leica DMI 6000 B (Leica Microsystems, USA). Percentage TUNEL positive cardiomyocytes were calculated with total number of cells using ImageJ software. A similar procedure was also used for the heart cryosection.

### Immunofluorescence and confocal microscopy

Cells were fixed with 4% paraformaldehyde and permeabilized with 0.1% Triton-X-100. Cells were washed with 1X PBS and incubated with 5% BSA for blocking. After blocking, all cells were incubated overnight with YTHDF3 primary antibody (#PA5-97034, Thermo Scientific, USA). The following day, cells were incubated with Alexa-Flour labeled secondary antibody (Thermo Scientific, USA). DAPI was used for nucleus staining. Images were taken with 20L×Lobjective using Leica DMI 6000 B (Leica Microsystems, USA) to get the localization of YTHDF3.

For Z-Staking imaging, neonatal rat cardiomyocytes were seeded on laminin-coated coverslips in 24 well plates and stained with α-sarcomeric actinin and YTHDF3 primary antibodies. Coverslips containing cardiomyocytes were mounted with 80% glycerol onto the glass slide and imaged in a confocal microscope (Olympus). For Z-staking, 10 images were captured for each cell having a distance of 1 μm.

### Animal experiments

For all the animal experiments C57BL/6 and BALB/c mice were provided by the National Laboratory Animal Centre, CSIR-CDRI, Lucknow, India. Mice from both strains were injected with 1.8*10^12^ AAV9 viral particles intravenously. Cardiac function was measured by echocardiography using Vevo 1100 (Fujifilm Visual Sonics, USA) under 2–3% isoflurane given by inhalation. Echocardiography was done at 25 days post viral injection. Mice were euthanized by cervical dislocation after anesthesia with isoflurane (3%), and the heart was taken out for further molecular, histological, morphological, ultrastructural, and molecular assays. The right tibia bone is isolated for length measurement. AAV9-scramble shRNA was used as a control. All the animal experiments were approved by the local IAEC (Institutional Animal Ethics Committee) committee at CSIR-Central Drug Research Institute following the guidelines of the Committee for the Purpose of Control and Supervision of Experiments on Animals (CPCSEA), New Delhi, Government of India. All the animal procedures performed conform to the guidelines from Directive 2010/63/EU of the European Parliament on the protection of animals used for scientific purposes. The heart weight and wet weight of the lung were normalized to the tibia length.

### YTHDF3 Pull down

H9c2 cells were seeded in three 150mm culture dishes (12 million cells per dish). Cells were washed with 1XPBS, trypsinized, and harvested cells were treated with 2mM DSP (Thermo Scientific, USA) for 15 min in a CO2 incubator, and the reaction was stopped using 1M Tris, pH 7.5 for 10 minutes. Cells were again washed with 1X PBS and subjected to nuclear-cytoplasmic fractionations as described above. One ml of 1XNP40 Cell lysis buffer was added to the nuclear pellet on ice for 15 minutes and sonicated for 15 minutes (10 seconds on and 5 seconds off phase with 30% amplitude) and further centrifuged at 12000*g, and nucleoplasm was collected for YTHDF pull down. YTHDF3 pull-down is carried out using Dynabeads antibody coupling kit (Thermo Scientific, USA). In brief, 10mg dynabeads were washed with C1 and C2 buffer, and incubated at 37°C overnight with 10μg of antibody (YTHDF3 and IgG control). The following day, beads were collected and washed with HB and LB buffer (2 times each) using magnetic stands and stored at 4°C for later use. Already collected 1 ml nucleoplasm was distributed to IgG and YTHDF3 antibody-bound beads (500μl each) and incubated for 4 hours at 4°C on a rotator. Beads were washed sequentially with IP wash buffer, high salt buffer, and phosphate wash buffer 2 times using an IP rotator. Finally, beads were dissolved in 1ml of PNK buffer. 30% of beads were used to confirm the pull-down by western blotting, and 70% were used for LC-MS/MS analysis.

### LC-MS/MS

Vproteomics, Valerian Chem Private Limited, India, performed LC-MS/MS analysis. Beads were separated from the PNK buffer using a magnetic stand, and bound proteins were digested with trypsin for 16hours at 37°C. Digested peptides were first cleaned using a C18 silica cartridge and then dried. The dried peptide pellet was dissolved in a buffer with 2% acetonitrile and 0.1% formic acid. Ultimate 3000 RSLC nano system coupled with a Tribrid Orbitrap Eclipse (Thermo Fisher Scientific, USA) is used for mass spectrometric analysis. 1 μg of the sample was loaded on Acclaim PepMap 75 μmL×L2 cm C18 guard column (3 μm particle size). Elution of peptide was done with a 0–40% gradient of a buffer having 80% acetonitrile and 0.1% formic acid and separated on a 50 cm, 3 μm Easy-spray C18 column (Thermo Fisher Scientific, USA) at a flow rate of 300 nl/min and further injected for MS analysis. LC gradient running was done for 110 min. MS1 spectra were acquired in the Orbitrap (RL=L240 k; AGQ targetL=L400,000; Max ITL=L50 ms; RF LensL=L30%; mass rangeL=L375L−L1500 m/z; centroid data). Dynamic exclusion was employed for 10 seconds, excluding all charge states for a given precursor. MS 2 spectra were collected in the linear ion trap (rateL=LRapid; AGQ targetL=L30,000; MaxITL=L20 ms; NCEHCDL=L30%). Raw data Generated by Mass spectroscopy was analyzed with Proteome Discoverer (v2.2) software against the UniProt *Rattus norvagicus* proteome (UP000002494) database, and data was exported as an Excel file having the number of peptides detected, PEP score and coverage. Unique peptides were selected by comparing them with IgG-bound peptides. The quantitative composition of the proteome is generated using the Bionic-vis program. Protein ID and the number of peptides were used as input.

### Transmission electron microscopy

Heart tissue from AAV9-*Ythdf3* shRNA and AAV9-Scramble shRNA injected mice were explanted and chopped at the site of the left ventricle in small pieces (1mm^3^) for transmission electron microscopy. Transmission electron microscopy was performed using the method already described, with minor modifications [2]. Tissues were fixed in 2.5% glutaraldehyde and 0.4% formaldehyde in 0.1M phosphate buffer, pH 7.4. Post-fixation tissue was treated with 1% osmium tetroxide and followed by dehydration steps in an ascending series of ethanol, infiltration, and embedded in Spurr resin. Ultra sectioning (60-70nm) of Spurr embedded tissue sections was performed using Leica EM UC7 ultramicrotome (Wetzlar, Germany), picked up on 200 mesh copper grid, and dual staining was performed using uranyl acetate and lead citrate. Grids were observed under JEOL JEM1400 TEM. All the images of TEM were captured using a Galton Orius SC 200B CCD camera at 100kV with Galton digital micrograph software.

### RNA sequencing and analysis of alternative splicing

Global transcriptomics was performed at Innovate Life Science, India. RNA was isolated from AAV9-injected animals by trizol based classical method. Isolated RNA was quantified using a NanoDrop ND-100 spectrophotometer (NanoDrop Technologies, Wilmington, DE), and RNA quality was assessed with the Tapestation 4200^TM^ (Agilent Technologies, Santa Clara, USA). Quality and control passed RNA was processed for RNA-seq library generation using KAPA RNA HyperPrep kit protocol with RiboErase (Roche Molecular Systems, Pleasanton, USA). In brief, the Riboerase step was performed to deplete ribosomal RNA from total RNA. The rRNA-depleted RNA was fragmented into small fragments(200-300bp) and reverse transcribed into cDNA using KAPA’s enzymes mix. cDNA was subjected to adapter ligation followed by PCR amplification. Next, the library was again quantified to determine the concentration and fragment size distribution. Finally, the library was sequenced on Illumina Novaseq 6000 to generate 40M, 2x150bp reads/sample, and 75% of the sequenced bases were of Q30 values>90.

The quality of sequencing reads was assessed using FASTQC, and low-quality reads, and adapter sequences were removed. The reference genome and annotation files for *Mus musculus* (GRCm39) were retrieved from the NCBI genome database. A genome index was generated using HISAT2, followed by read alignment to the indexed genome. The aligned reads were stored as SAM files and converted to BAM format using SAMtools for further processing.

Alternative splicing analysis was conducted using rMATS (RNA-seq Multivariate Analysis of Transcript Splicing). The sorted BAM files from control and test samples were compiled into separate input files for rMATS. Differential alternative splicing events, including skipped exons (SE), mutually exclusive exons (MXE), alternative 5’ splice sites (A5SS), alternative 3’ splice sites (A3SS), and retained introns (RI), were identified. The analysis was conducted with a read length parameter of 150 bp, allowing variable read lengths. Statistically significant splicing events were filtered based on the false discovery rate (FDR<0.05) and inclusion level difference (δPSI±0.2) to ensure reliable and biologically meaningful results.

### Statistics

All the data analysis was performed with GraphPad Prism software. All the data is represented as mean±SEM. Different statistical tests (student t-test or ANOVA) were performed to evaluate significant differences. The calculated p-value is given in the figure legends.

## Results

### YTHDF3 knockdown in vitro induces cardiac atrophy and apoptosis

To study the function of *Ythdf3* in cardiomyocytes, we designed shRNA targeting *Ythdf3* and produced the AAV6 (Adeno-associated virus serotype 6) virus. We successfully silenced *Ythdf3* in isolated neonatal rat cardiomyocytes using AAV6-shRNA-*Ythdf3* at mRNA (**Figure 1A**) and protein (**Figure 1B**). Knockdown of *Ythdf3* in cardiomyocytes led to a significant decline in cardiomyocyte size, as evidenced by reduced cardiomyocyte area (**Figure 1C-D**). Furthermore, *Ythdf3* knockdown significantly increased cardiomyocyte death, as measured by Tunel staining (**Figure 1E-F**). To confirm that the atrophic and apoptotic effects directly result from its knockdown, we simultaneously overexpressed *Ythdf3* in neonatal rat cardiomyocytes together with its knockdown. *Ythdf3* cDNA was amplified from mouse heart and cloned in an AAV vector under a cTnT promoter for overexpression. Overexpression was confirmed in neonatal rat cardiomyocytes (**Figure 1G**). Overexpression of *Ythdf3* rescued knockdown-induced cardiac atrophy (**Figure 1H-I**) and apoptosis (**Figure 1J-K**). To further confirm the deleterious effects of *Ythdf3*, we knockdown *Ythdf3* in a rat cardiomyocyte cell line H9c2 using lentivirus (**Supplementary Figure 1A-B**). Knockdown of *Ythdf3* significantly induced the death of H9c2 cells as measured by MTT assay (**Supplementary Figure 1C**) and Tunel staining (**Supplementary Figure 1D-E**). Next, to confirm that cell death induced by *Ythdf3* shRNA is not due to any off-target effect of shRNA itself, we further designed two additional shRNAs targeting *Ythdf3* and tested its effect in H9c2 cells. Lentivirus-mediated delivery of shRNA 2&3 in H9c2 cells also successfully silenced *Ythdf3* (**Supplementary Figure 1F**). Knockdown of *Ythdf3* by shRNAs 2&3 also led to a significant decline in cell survival (**Supplementary Figure 1G**). These results confirm that lower levels of *Ythdf3* lead to cardiomyocyte atrophy and death.

**Fig. 1.**
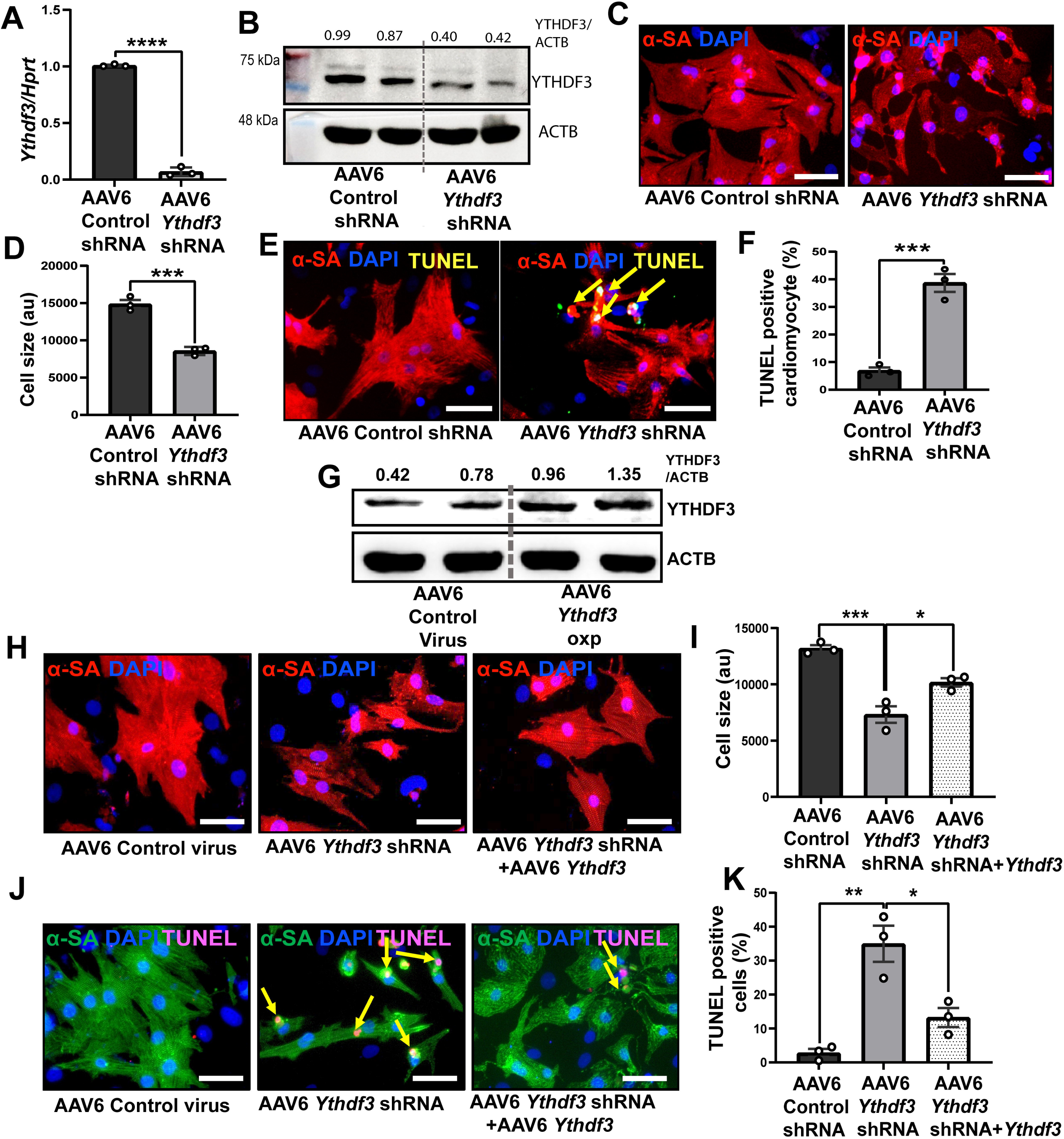
AAV6 mediated knockdown of *Ythdf3* induces cardiomyocyte atrophy and apoptosis in primary cardiomyocytes. (**A-B)** AAV6 mediated knockdown of *Ythdf3* in neonatal rat cardiomyocytes at mRNA level (**A**) and at protein level (**B**) (*n*=3, Student t-test, unpaired (two-tailed) ****p<0.0001 (**A**)). (**C-D)** Cell size of neonatal rat cardiomyocytes transduced with AAV6 Control shRNA and *Ythdf3* shRNA, where **C** shows representative image (n=3, Student t-test, unpaired (two-tailed) \*\*\**p*=0.0006). (**E-F)** TUNEL staining for apoptosis in neonatal rat cardiomyocytes transduced with AAV6 Control shRNA and *Ythdf3* shRNA (Yellow arrow shows TUNEL positive cells), where **E** shows representative image, *n*=3, Student t-test, unpaired (two-tailed) \*\*\**p*=0. 0008. (**G**) Western blot showing YTHDF3 levels in neonatal rat cardiomyocytes transduced with AAV6 cTnT Control and AAV6 cTNT *Ythdf3* virus at MOI of 5*10^3^. (**H-I)** The cell size of neonatal rat cardiomyocytes upon *Ythdf3* knockdown and simultaneous *Ythdf3* overexpression, **H** shows representative image (*n*=3, one-way ANOVA with Tukey’s multiple comparisons test \*\*\**p*=0.0004, \**p*=0.017). **(J-K)** TUNEL staining in neonatal rat cardiomyocytes upon *Ythdf3* knockdown and simultaneous *Ythdf3* overexpression where J shows representative image, *n*=3, one-way ANOVA with Tukey’s multiple comparisons test \*\**p*=0.0017, \**p*=0.011. α-SA - alpha sarcomeric actinin. au - arbitrary unit. Scale bar represents 50μm.

Similar to *Ythdf3* knockdown, doxorubicin, a well-known cardio-toxic agent, also induces cardiomyocyte death and atrophy. Hence, we wanted to study whether doxorubicin alters *Ythdf3* levels in cardiomyocytes and thus mediates its cardiotoxic effects. We found a significant decline in *Ythdf3* protein levels in H9c2 cells treated with doxorubicin (**Figure 2A-B**). Next, to confirm *Ythdf3* regulation in response to doxorubicin, we treated adult BALB/c male mice with 5mg/kg doxorubicin once a week for four weeks **(Figure 2C)**. Doxorubicin treatment led to cardiac atrophy (**Figure 2D**), which is evident by a significant decline in heart-weight to tibia-length ratio (**Figure 2E**) and impaired cardiac function (**Figure 2F**). Significant downregulation of *Ythdf3* was observed in hearts exposed to doxorubicin, confirming the *in vitro* data (**Figure 2G**). Furthermore, overexpression of *Ythdf3* rescued doxorubicin-induced cardiomyocyte atrophy (**Figure 2H-I**) and death (**Figure 2J-K**) in neonatal rat cardiomyocyte, thus showing it as an essential regulator of doxorubicin-mediated cardiomyocyte atrophy and death.

**Fig. 2.**
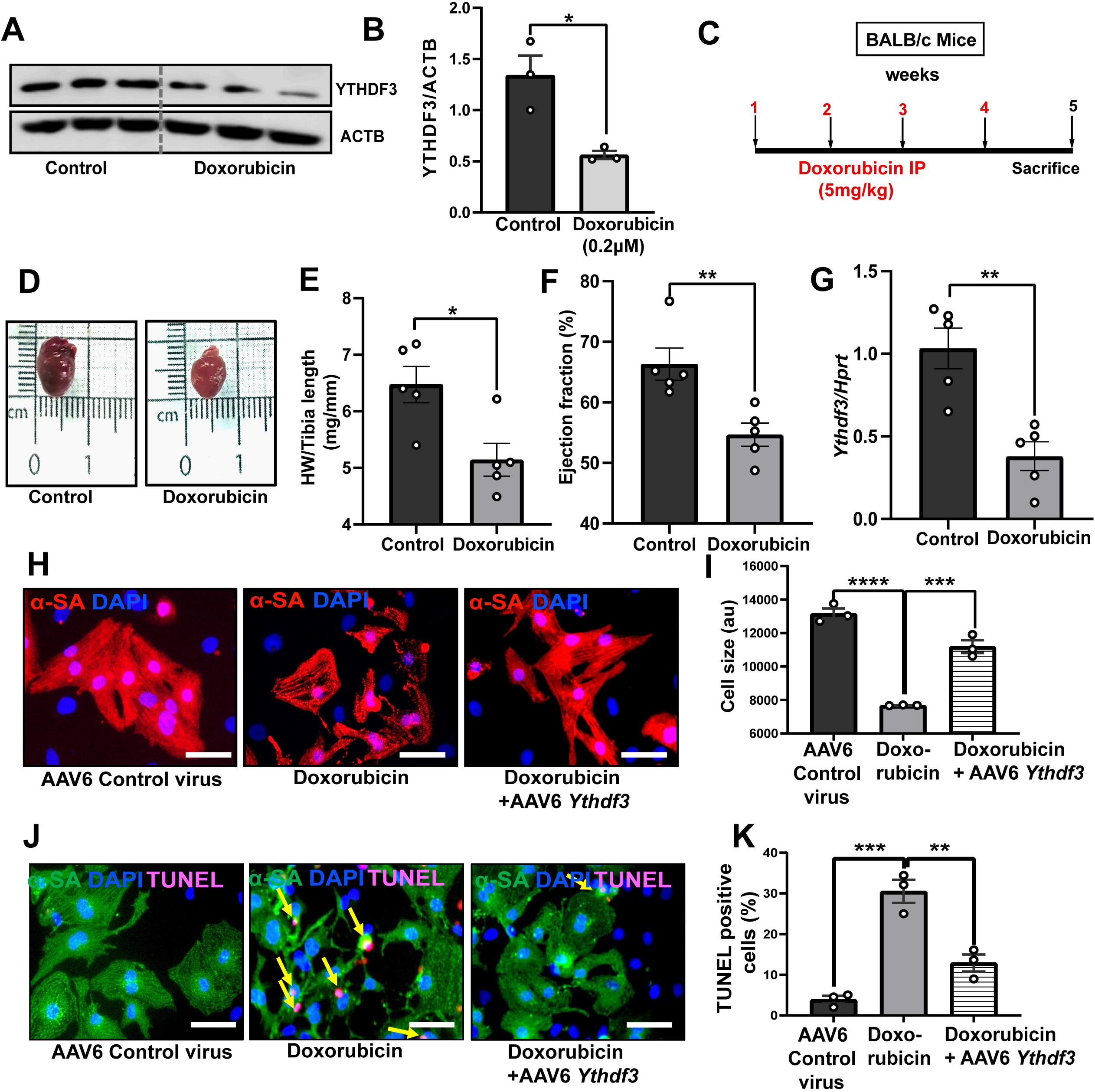
Ythdf3 prevents doxorubicin-induced atrophy and apoptosis. **(A-B)** Western blot showing decreased protein expression of YTHDF3 in doxorubicin treated H9c2 cells (n=3) Student t-test, unpaired (two tailed) \**p*=0. 0171. **C** Schematic representation of doxorubicin induced cardiac atrophic mouse model. **(D)** Gross morphology of control and doxorubicin treated heart. **(E)** Heart weight to tibia length ratio control (*n*=5) doxorubicin (*n*=5) Student t-test, unpaired (two tailed) \**p*<0.0153. **(F)** Ejection fraction Student t-test, unpaired (two tailed)\*\**p*=0.0075. **(G)** mRNA expression of *Yhdf3* in control and doxorubicin treated mouse heart, Student t-test, unpaired (two tailed) \*\**p*=0.0025. (**H-I)** Cell size rescue experiment in doxorubicin treated NRCMs with AAV6 mediated *Ythdf3* overexpression, **H** shows representative image(*n*=3) one-way ANOVA with Tukey’s multiple comparisons test \*\*\*\**p*=0.0001, \*\*\**p*= 0.0003. **(J-K)** TUNEL rescue experiment in doxorubicin treated NRCMs with AAV6 mediated *Ythdf3* overexpression where **K** shows representative image (*n*=3) one-way ANOVA with Tukey’s multiple comparisons test \*\*\**p*=0.0003, \*\**p*=0.0025. α-SA - alpha sarcomeric actinin, au -arbitrary unit. The scale bar represents 50µm.

### Knockdown of *Ythdf3* in murine heart induces cardiac cachexia

Next, to further confirm the *in vitro* effects of *Ythdf3* depletion on cardiomyocytes, we knockdown *Ythdf3* in adult murine C57BL6 hearts using a similar shRNA strategy delivered by AAV9. AAV9 virus shows cardio-tropism and is used for specific gene transfer to cardiomyocytes in the heart [16, 21]. AAV9-mediated delivery of shRNA significantly decreased the levels of *Ythdf3* in the heart (**Figure 3A**). *Ythdf3* is a ubiquitous protein that is expressed by non-cardiomyocyte cells, too. We isolated adult cardiomyocytes from AAV9-*Ythdf3*-shRNA-treated hearts and control to confirm the knockdown in cardiomyocytes. *Ythdf3* levels were confirmed to be declined in cardiomyocytes treated with AAV9-*Ythdf3*-shRNA, confirming cardio-tropism (**Figure 3B**). Knockdown of *Ythdf3* led to a significant decline in the body weight of mice compared to scramble shRNA (**Figure 3C**). *Ythdf3* knockdown led to a substantial decline in heart mass, confirming cardiac atrophy (**Figure 3D-E**). Furthermore, echocardiography measured left ventricular mass significantly decreased upon *Ythdf3* knockdown (**Figure 3F**). To further confirm the cardiac atrophy at the cardiomyocyte level, wheat-germ agglutinin staining measured the cardiomyocyte area in a cross-sectional view. We found a significant decline in cardiomyocyte cross-sectional area upon knockdown of *Ythdf3* (**Figure 3G-H**). To further confirm the effect on cardiomyocytes, we isolated adult cardiomyocytes from *Ythdf3* knockdown hearts and measured their surface area. Cardiomyocytes from *Ythdf3* depleted hearts exhibited significantly decreased total surface area, length, and width compared to control, confirming cardiomyocyte atrophy (**Figure 3I-L**). Ultrastructure visualization by electron microscopy found a decline in the density of myofibres, further confirming atrophy (**Figure 3M**). Additionally, the knockdown of *Ythdf3* in the heart resulted in significantly increased Tunel-positive nuclei (**Figure 3N-O**), and increased cleaved caspase-3 expression (**Figure 3P-Q**), depicting increased cardiomyocyte death. *Ythdf3* knockdown lungs have increased water content, showing signs of congestive heart failure (**Figure 3R**). In line with this, cardiac function measured by echocardiography also significantly decreased in *Ythdf3* knockdown hearts (**Figure 3S**). These results are in line with the *in vitro* data confirming cardiac cachexia upon knockdown of *Ythdf3*.

**Fig. 3.**
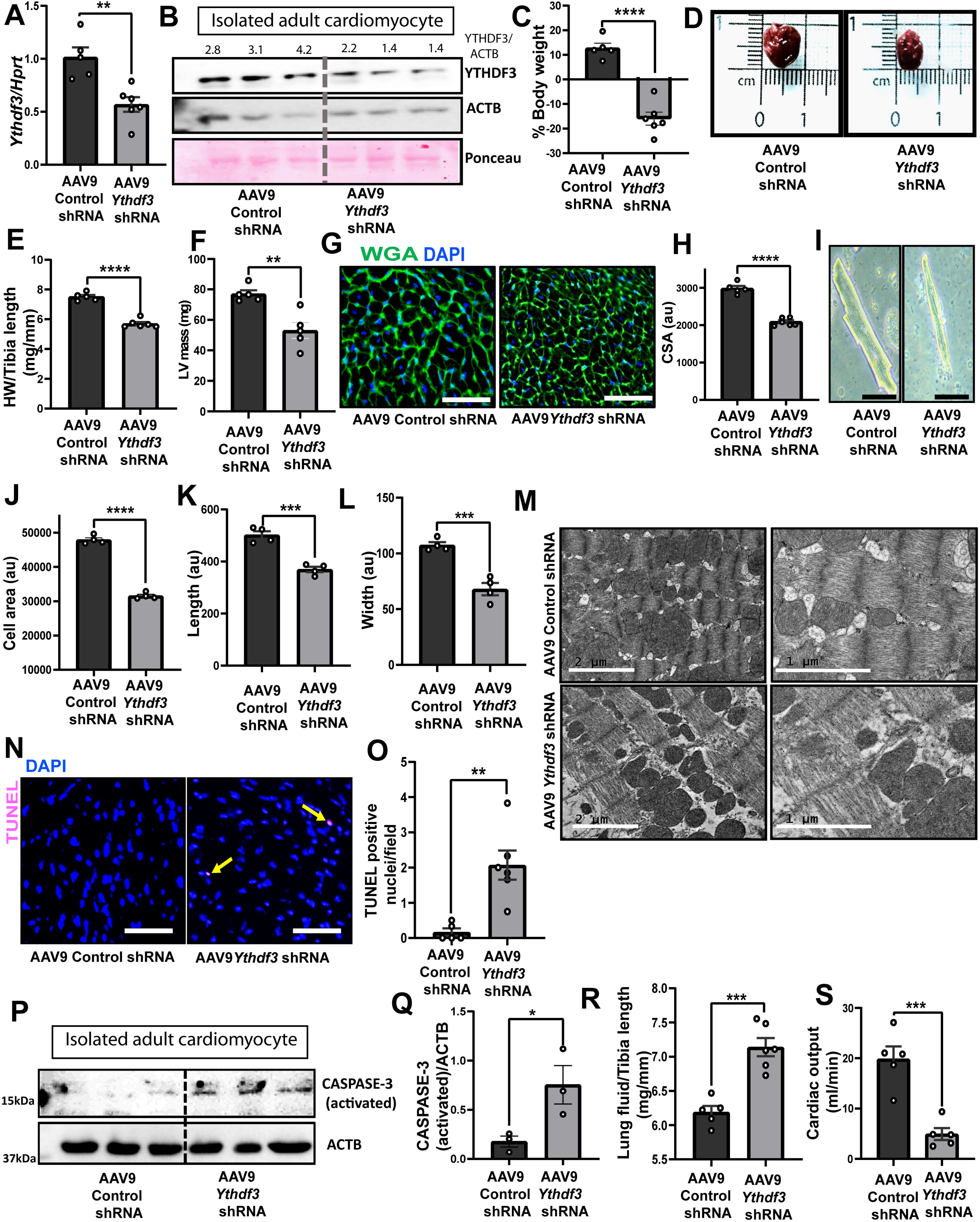
AAV9 mediated knockdown of *Ythdf3* leads to cardiac cachexia in C57BL/6 mice. **(A)** Expression levels of *Ythdf3* in total heart of AAV9 Control and *Ythdf3* shRNA injected mice, Control (n= 5) and *Ythdf3* knockdown (n=6) Student t-test, unpaired (two-tailed) \*\**p*= 0.0032. **(B)** Western blot showing *Ythdf3* at protein level in adult cardiomyocytes isolated from AAV9 Control (*n*=3) and *Ythdf3* shRNA (*n*=3) injected animals. (**C)** Percentage body weight change of animals injected with Control (*n*=5) and *Ythdf3* shRNA (n=6), Student t-test, unpaired (two-tailed) \*\*\*\**p*< <0.0001. **(D)** Gross morphology of Control and *Ythdf3* depleted hearts. **(E)** Heart weight to tibia length ratio of Control (n=5) and *Ythdf3* knockdown (n=6) hearts, Student t-test, unpaired (two-tailed) \*\*\*\**p*<0.0001. (**F)** Left ventricular mass analyzed by echocardiography Control (n=5) and *Ythdf3* knockdown (n=5) Student t-test, unpaired (two-tailed) \*\**p*=0.0031. **(G-H)** Cross-sectional cardiomyocyte area of heart stained with WGA and DAPI where **(G)** shows representative image, Student t-test, unpaired (two-tailed) \*\*\*\**p*<0.0001. The scale bar represents 50µm. **(I-L)** Cell size measurements of adult cardiomyocyte isolated from Control and *Ythdf3* knockdown mice hearts where **I** shows representative image, cell area (**J**), cardiomyocyte length (**K**), cardiomyocyte width (**L**), Control (*n*=4) *Ythdf3* knockdown (*n*=4) Student t-test, unpaired (two-tailed) \*\*\*\**p*<0.0001 (**J**), \*\*\**p*=0.0003 (**K**), \*\*\**p*=0.0007 (**L**). The scale bar represents 50 µm. (**M)** Transmission electron microscopy image showing damaged sarcomere in *Ythdf3* knockdown heart. The scale bar represents 1µm and 2 µm. **(N-O)** TUNEL staining of Control (n= 5) and *Ythdf3* depleted hearts (n=6) where **N** shows representative image, Student t-test, unpaired (two-tailed) \*\**p*=0.0028. (**P**-**Q**) Expression level of cleaved caspase-3 upon Ythdf3 knockdown and control, (n=3) Student t-test, unpaired (two-tailed) \**p*=0.0462. **(R)** Lung fluid content to tibia length ratio Control (n=5) and *Ythdf3* knockdown (n=6), Student t-test, unpaired (two-tailed) \*\*\**p*=0.0006. **(S)** Cardiac output measured by echocardiography Control (n=5) and *Ythdf3* knockdown (n=5), Student t-test, unpaired (two-tailed) \*\*\**p*=0.0006.

To further confirm that *Ythdf3* knockdown leads to cardiac cachexia, we tested its knockdown effects in an additional mice strain, BALB/C. Adult BALB/C mice treated with AAV9-*Ythdf3*-shRNA confirmed significant downregulation of *Ythdf3* in the heart at mRNA (**Supplementary Figure 2A**) and protein levels (**Supplementary Figure 2B-C**). Similar to C57Bl/6, knockdown of *Ythdf3* in BALB/C mice led to a significant decline in body weight (**Supplementary Figure 2D**), heart weight-tibia-length ratio (**Supplementary Figure 2E**), left ventricular mass (**Supplementary Figure 2F**) measured by echocardiography and cardiomyocyte cross-sectional area (**Supplementary Figure 2G-H**). Similarly, Tunel-positive nuclei were also increased in *Ythdf3* knockdown hearts (**Supplementary Figure 2I-J**). Knockdown of *Ythdf3* led to accumulation of fluid in the lungs (**Supplementary Figure 2K**) and functional impairment (**Supplementary Figure 2L**). These results with two independent strains confirm that *Ythdf3* knockdown leads to cardiac cachexia and impaired function.

### Nuclear localization of YTHDF3 in cardiomyocytes

The function of RNA-binding proteins depends upon their localization. YTHDF3 is known to localize mainly in the cytoplasm; however, its nuclear localization has also been reported [6, 25]. Hence, to explore the downstream functions of YTHDF3, its cellular localization in cardiomyocytes is a prerequisite. We found primarily nuclear localization of YTHDF3 in isolated neonatal rat cardiomyocytes, rat cardiomyocyte cell line H9c2, neonatal mouse cardiomyocytes, and adult mouse cardiomyocytes by immunofluorescence staining (**Figure 4A**). To further confirm nuclear localization, we performed imaging at different focal planes along the z-axis in neonatal rat cardiomyocytes. Z-stacking confirmed nuclear localization of YTHDF3 (**Figure 4B**).

**Fig. 4.**
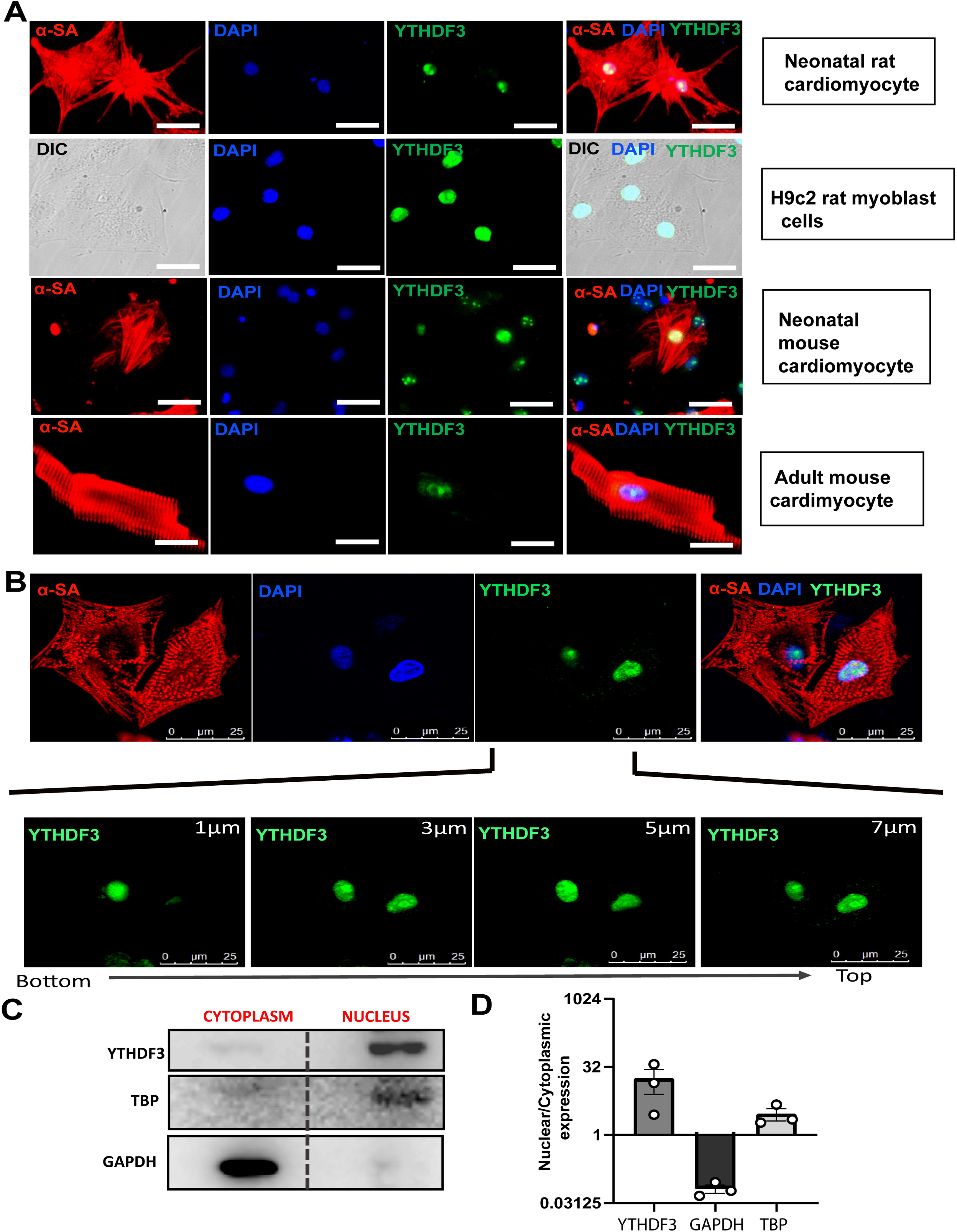
YTHDF3 uniquely localizes to the nucleus in cardiomyocytes. **(A)** Representative image showing YTHDF3 nuclear localization in neonatal rat cardiomyocytes, H9c2 myoblast cells, neonatal mouse cardiomyocytes, and adult mouse cardiomyocytes. The scale bar represents 50µm. **(B)** Confocal microscope z-stacks projections showing the localization patterns of YTHDF3 at different depth of isolated neonatal rat cardiomyocytes. The scale bar represents 25µm. (**C-D)** Western blot showing the nucleus-cytoplasmic ratio of proteins in H9c2 cells.

Furthermore, we separated the cytoplasmic and nucleoplasmic proteome of H9c2 cells and found YTHDF3 highly enriched in nucleoplasm (**Figure 4C-D**). TBP and GAPDH were used as controls for nucleoplasmic and cytoplasmic fractions, respectively (**Figure 4C-D**). Nuclear localization was further confirmed in isolated adult mouse cardiomyocytes, neonatal rat cardiomyocytes, and rat cardiomyocyte cell line H9c2 using a different antibody, ruling out any off-target effect due to misrecognition by the antibody (**Supplementary Figure 3A**). Additionally, we checked the localization of YTHDF3 in other cell lines. Human hepatocyte line HePG2 and human cervical cancer Hela cell line showed cytoplasmic localization, while mouse fibroblast cell line L929 showed nuclear localization (**Supplementary Figure 3B**). These results confirm cell-dependent localization of YTHDF3 to cytoplasm or nucleus.

### YTHDF3 regulates alternative splicing in cardiomyocyte

Nuclear-localized RNA-binding proteins primarily regulate splicing and transport of RNAs. Protein interactome provides an important clue to understanding the function of a protein. Therefore, we designed an experiment with a pull-down of YTHDF3 followed by LC-MS/MS analysis (**Figure 5A**). We performed immunoprecipitation of YTHDF3 and validated it by western blotting (**Figure 5B**). Next, to delineate the functions of YTHDF3 in cardiomyocytes, we subjected the immunoprecipitated protein complex by YTHDF3 antibody and IgG control to LC-MS/MS analysis. We identified 108 unique proteins in the YTHDF3 interactome complex compared to IgG. To understand the function of interacting proteins, we performed a quantitative composition of proteins focusing on protein function using Proteomaps. We found a significant overrepresentation of terms related to RNA processing, including ribosome, RNA transport, and spliceosome which has the highest enrichment (**Figure 5C**). Among the *Ythdf3* interacting proteins involved in spliceosomes, DDX5 and HNRNPU have been shown to regulate alternative splicing in the heart and interact with each other [13, 22]. We further confirmed the interaction of YTHDF3 with DDX5 and HNRNPU by co-immunoprecipitation and western blot (**Figure 5D**). These results suggest that YTHDF3 may regulate alternative splicing events in the heart.

**Fig. 5.**
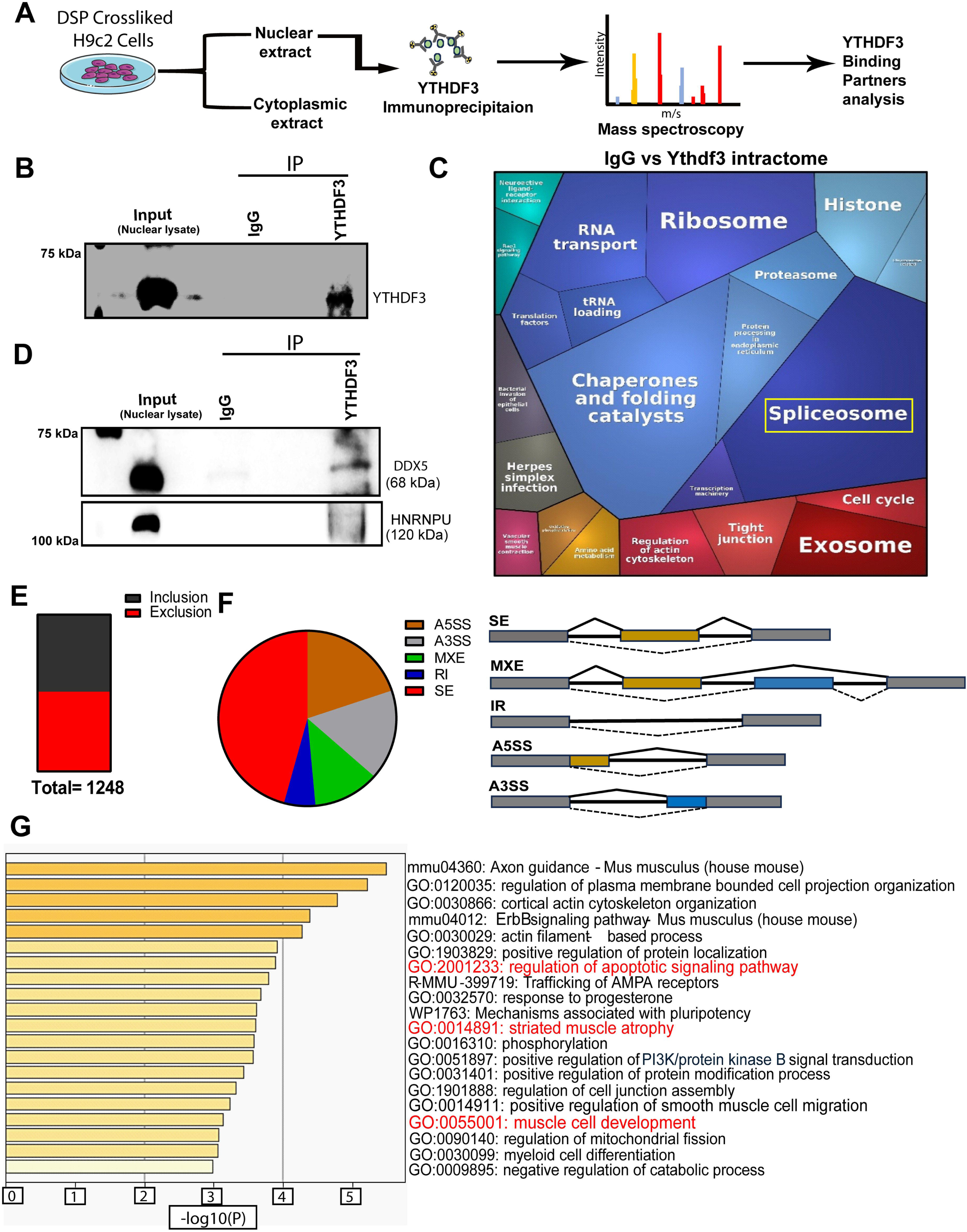
YTHDF3 majorly interacts with RNA splicing complex and regulates alternative splicing. **(A)** Schematic representation for immunoprecipitation–mass spectrometry (IP-MS) analysis. **(B)** Western blot showing *Ythdf3* immunoprecipitation **(C)** Proteo-map representing YTHDF3 bound proteins interactome analysis. **(D)** Western blot of DDX5 and HNRNPU from YTHDF3 and IgG control pull-down samples. **(E)** The distribution of inclusion and exclusion among alternatively spliced events. (**F**) Pie-chart showing the distribution of alternatively spliced events, SE=exon skipping, MXE= mutually exclusive exons, RI= intron retention, A5SS=alternative 5’splice site, A3SS= alternative 3’splice site. **(G)** Gene ontology (GO) term analysis of differentially spliced skipped exon genes upon *Ythdf3* knockdown.

Next, we performed RNA-seq from shRNA-control and shRNA-*Ythdf3* hearts to investigate *Ythdf3*-regulated alternative splicing. The rMATS software was used to study the differential splicing events with a cut-off of ΔPSI≥±0.2 and FDRL0.05. A total of 1248 events (**Supplementary Table 3**) were found to be differentially spliced with almost equal distribution for inclusion and exclusion events (**Figure 5E**). The differential spliced events included skipped exons (SE), alternative 5’ splice site (A5SS), alternative 3’ splice site (A3SS), mutually exclusive exons (MXE), and retained introns (**Figure 5F**). Skipped exons (SE) were the highest splicing event to be altered, covering more than 45% of the 1248 events (**Figure 5F**). Gene ontology enrichment analysis of SE genes revealed significant enrichment of terms related to apoptosis, muscle atrophy, and muscle cell development, confirming the phenotype of *Ythdf3*-shRNA hearts (**Figure 5G**). These data show that *Ythdf3* regulates alternative splicing events in the hearts.

### YTHDF3 regulates CaMKIIδ alternative splicing

YTHDF3 interacting partners DDX5 and HNRNPU have been shown to regulate alternative splicing of CaMKIIδ in the heart [13, 22]. Hence, we wanted to check whether YTHDF3 regulates alternative splicing of CaMKIIδ. CaMKIIδ forms multiple isoforms, CaMKIIδA, CaMKIIδB, CaMKIIδC, and CaMKIIδ9 isoforms, formed due to alternative splicing of exons 14, 15 and 16, are expressed in the heart [13, 22]. A schematic showing exons present in each isoform is shown in **Figure 6A**. We designed a primer pair binding in the constitutive exons 13 and 18 to amplify all isoforms simultaneously (**Figure 6B**). RNA was extracted from the isolated adult cardiomyocytes of BALB/C and C57BL6 mice heart injected with shRNA targeting *Ythdf3* and scramble control. We found increased levels of CaMKIIδA and CaMKIIδC isoform upon knockdown of *Ythdf3* in both BALB/c (**Figure 6C**) and C57BL6 (**Figure 6D**) cardiomyocytes. Levels of CaMKIIδ9 isoform were found to be decreased upon *Ythdf3* knockdown (**Figure 6C-D**). CaMKIIδB isoform was very weakly expressed in both the cardiomyocyte samples (**Supplementary Figure 4**). Similar to our data, DDX5 and

**Fig. 6.**
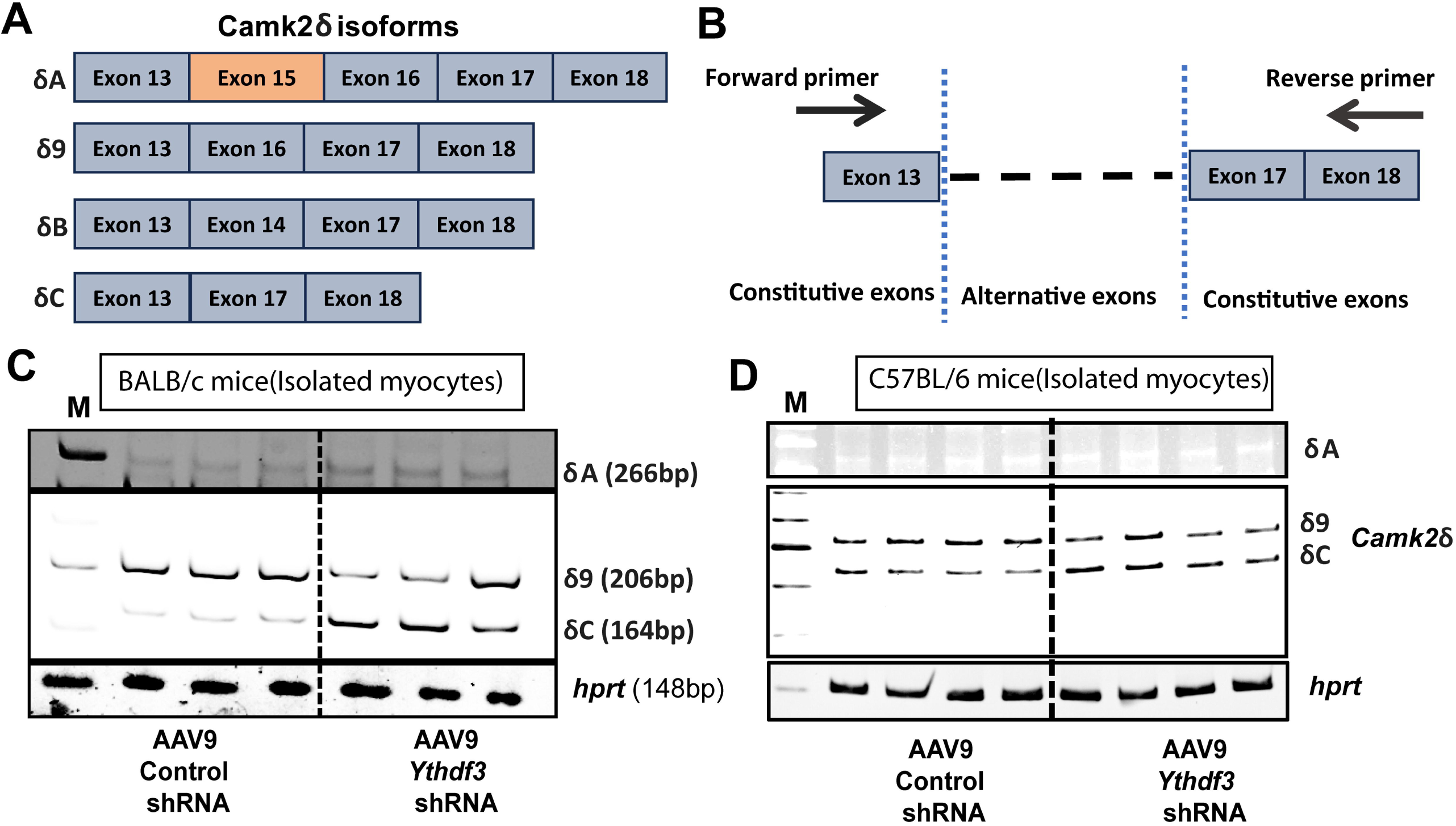
YTHDF3 regulates alternative splicing of CaMKIIδ in the mouse heart. **(A)** Illustration of different spliced isoforms of CaMKIIδ gene expressed in the heart: (δA, δ9, δB, and δC). (**B**) The image shows forward and reverse primer locations used to amplify all isoforms. **(C-D)** DNA-PAGE gel showing expression of different CaMKIIδ isoforms from isolated adult cardiomyocytes of Control and *Ythdf3* knockdown hearts from BALB/c (**C**) and C57BL/6 (**D**) mice.

HNRNPU deficient hearts also show increased levels of CaMKIIδA and CaMKIIδC isoforms while decline in CaMKIIδ9 [13, 22]. These results show that YTHDF3 regulates CaMKIIδ alternative splicing by interacting with DDX5 and HNRNPU.

## Discussion

M^6^A modification of RNAs adds a layer of post-transcriptional regulation to the cardiac transcriptome. Here, we have shown that the knockdown of m6A reader protein YTHDF3 in cardiomyocytes led to cardiac cachexia marked by atrophy and cardiomyocyte death, highlighting its indispensable role in the heart. Doxorubicin, a cardiotoxic agent known to induce cardiac cachexia, downregulated the expression of *Ythdf3* in vitro and in vivo, and its overexpression in isolated neonatal rat cardiomyocytes rescued the doxorubicin-induced atrophy and death showing *Ythdf3* as an essential regulator of cardiac cachexia.

YTHDF family members, including YTHDF3, have been mainly shown to localize in cytoplasm, bind 3’UTR of mRNAs, and regulate their stability and translation. Here, we have demonstrated that YTHDF3 localizes to the nucleus in cardiomyocytes, which is unique to YTHDF3 among its family members. YTHDF3 interacts with splicing regulators like DDX5 and HNRNPU in the nucleus and regulates the cardiac splicing program. Similar to DDX5 and HNRNPU, we found that *Ythdf3* regulates alternative splicing of CaMKIIδ. Interestingly, deletion of both DDX5 and HNRNPU also results in death and loss of cardiomyocytes [13, 22].

One limitation of our study is using a universal U6 promoter, which may affect other organs. However, similar *in vitro* and *in vivo* results support our study results as a direct consequence of knockdown in the heart rather than the indirect effect of other organs.

CaMKII is well known to regulate cell death in the heart, and its inhibition improves cardiac function and promotes cardiomyocyte survival in different cardiovascular disease models [8]. Overexpression of CaMKIIδC isoform has been shown to induce cardiomyocyte cell death, while its dominant negative variant prevents cell death in neonatal rat cardiomyocytes treated with H2O2 or acidic buffer [24]. We have found increased levels of CaMKIIδC isoform upon *Ythdf3* knockdown, which may be the downstream effector responsible for cardiomyocyte death. Another isoform, CaMKIIδ9, has also been shown to promote cardiomyocyte death and lead to heart failure [23]. However, caution should be taken in assessing the function of each isoform by overexpression of a single isoform because CaMKII different isoforms heteromultimerize to form holoenzyme, and their localization and function depend upon the ratio of different isoforms [8]. Hence, overexpression of a single isoform may disturb the composition of the CaMKII holoenzyme and its function.

Recently, several studies have demonstrated the role of m^6^A reader protein YTHDF2 in cardiomyocyte death. YTHDF2 had been shown to inhibit hypoxia/reperfusion induced cell death by targeting BNIP3 in H9c2 cells [3]. Very recently, YTHDF2 has been shown to improve cardiac function and promote cardiomyocyte survival in a model of myocardial infarction [19]. YTHDF2 was found to inhibit ferroptosis by targeting NCOA4 mRNA [19].

## Abbreviation

RBPs: RNA-binding proteins
AAV: Adeno-associated Viruses
TUNEL: Terminal deoxynucleotidyl transferase dUTP nick end labeling
TEM: Transmission Electron Microscope
FDR: False Discovery Rate
PSI: Percent Splicing Index
mRNA: Messenger RNA
DAPI: 4′,6-diamidino-2-phenylindole
AS: Alternative Splicing
SE: Skipped Exon
LC-MS/MS: Liquid Chromatography with tandem mass spectrometry
CaMKII: calcium/calmodulin-dependent protein kinase II
m6A: N6-methyladenosine
PBS: Phosphate buffer saline
cTnT: cardiac Troponin T

## Acknowledgments

We want to thank the Director, CSIR-CDRI, Dr. Manoj Kumar Barthwal, Dr. Kumaravelu Jagavelu, Dr. Chandra Prakash Pandey, Dr. Sheeba Saji Samuel, Mr. Pankaj Kumar Shukla, SAIF facility, and Animal House, CSIR-CDRI, for their help with the instruments and resources.

## Author contribution

SKG and AG have developed the concept, designed the study, planned and performed experiments, analyzed results, and prepared the manuscript. SC and SK (Shailesh Kumar) performed RNA-seq analysis. RKS and KM performed the transmission electronmicroscopy. GE helped with H9c2 cell experiments. RK performed in vivo doxorubicin experiment. SS and SK helped with isolation of neonatal rat cardiomyocytes and adult cardiomyocytes. SP helped in cloning and cell size experiments.

---

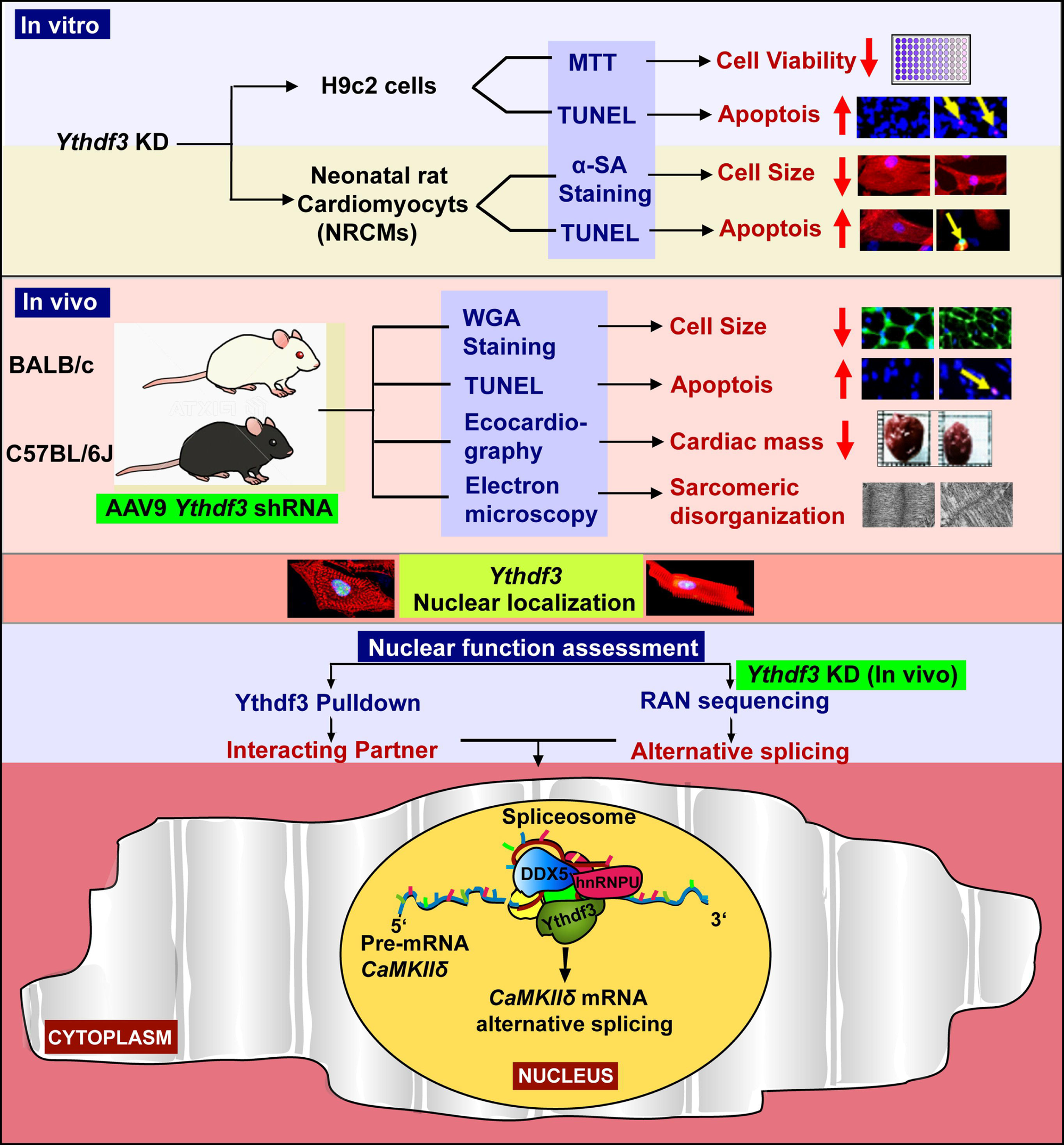

