## Supplementary material for "M^6^A reader protein YTHDF3 regulates cardiomyocyte death and atrophy by modulating the alternative splicing program": supplementary_ythdf3_manuscript f.pdf

#### Supplementary Fig. 1

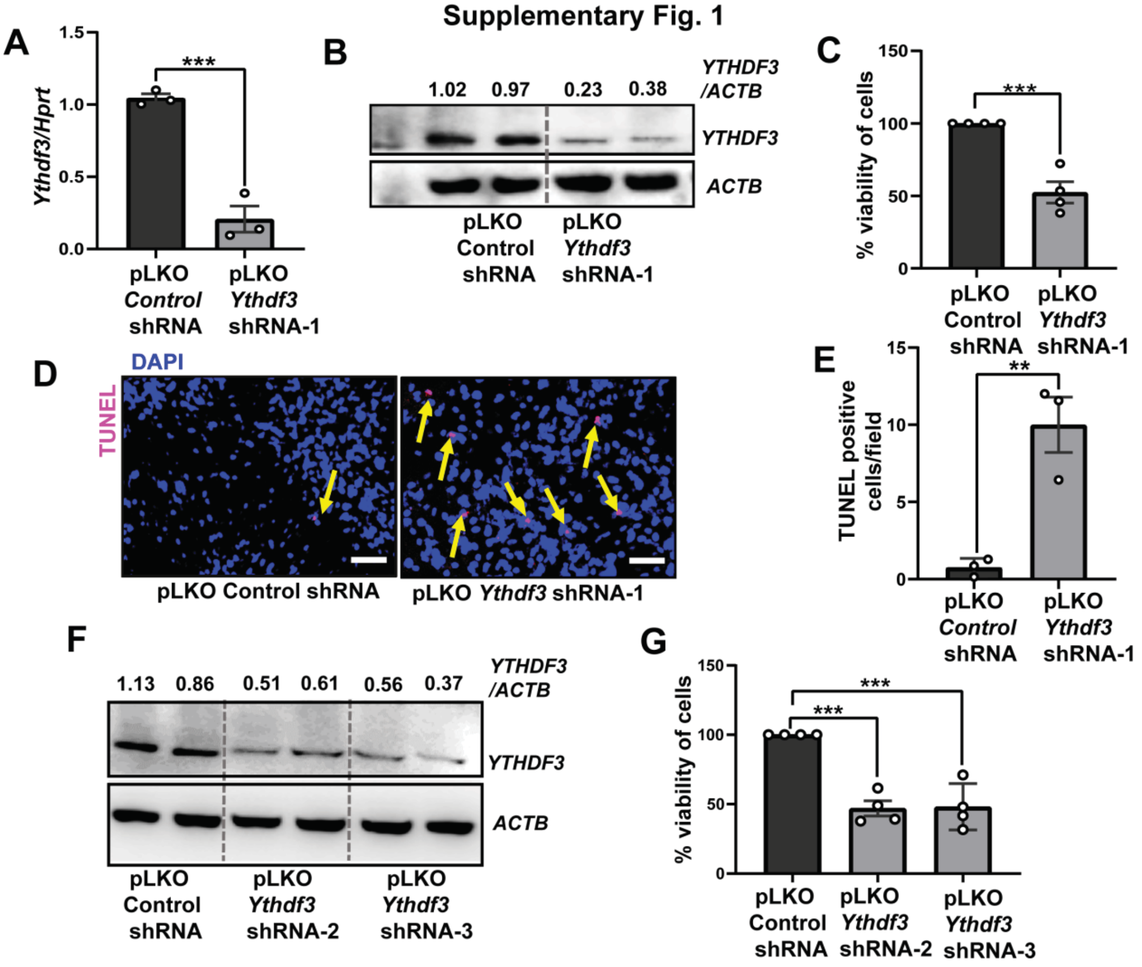


**Suplementary Fig 1. Lentiviral-mediated knockdown of *Ythdf3* leads to apoptosis in H9c2 cells.** **(A)** Expression level of *Ythdf3* in H9c2cells transduced with pLKO Control and *Ythdf3* shRNA-1 (*n*=3, Student t-test, unpaired (two-tailed) ***p=0.0009). **(B)** Western blot showing *Ythdf3* protein levels after transduction with pLKO Control and *Ythdf3* shRNA in H9c2 cells. **(C)** MTT assay for Viability (%) of H9c2 cells after *Ythdf3* knockdown (n=4, Student t-test, unpaired (two-tailed) ****p*<0.0006). **(D-E)** TUNEL staining for apoptosis in *Ythdf3* knockdown cells where **D** shows representative image (*n*=3, Student t-test, unpaired (two-tailed) **p=0.0071). **(F)** Western blot showing *Ythdf3* inhibition in H9c2 cells with pLKO *Ythdf3* shRNA 2 and 3 normalized with ACTB. **(G)** Percentage viability of cells after transduction with *Ythdf3* shRNA 2 and 3 (*n*=3, one-way ANOVA with Tukey's multiple comparisons test ****p*=0.0003, ****p*=0.0003). The scale bar represents 50μm.

Supplementary Fig. 2

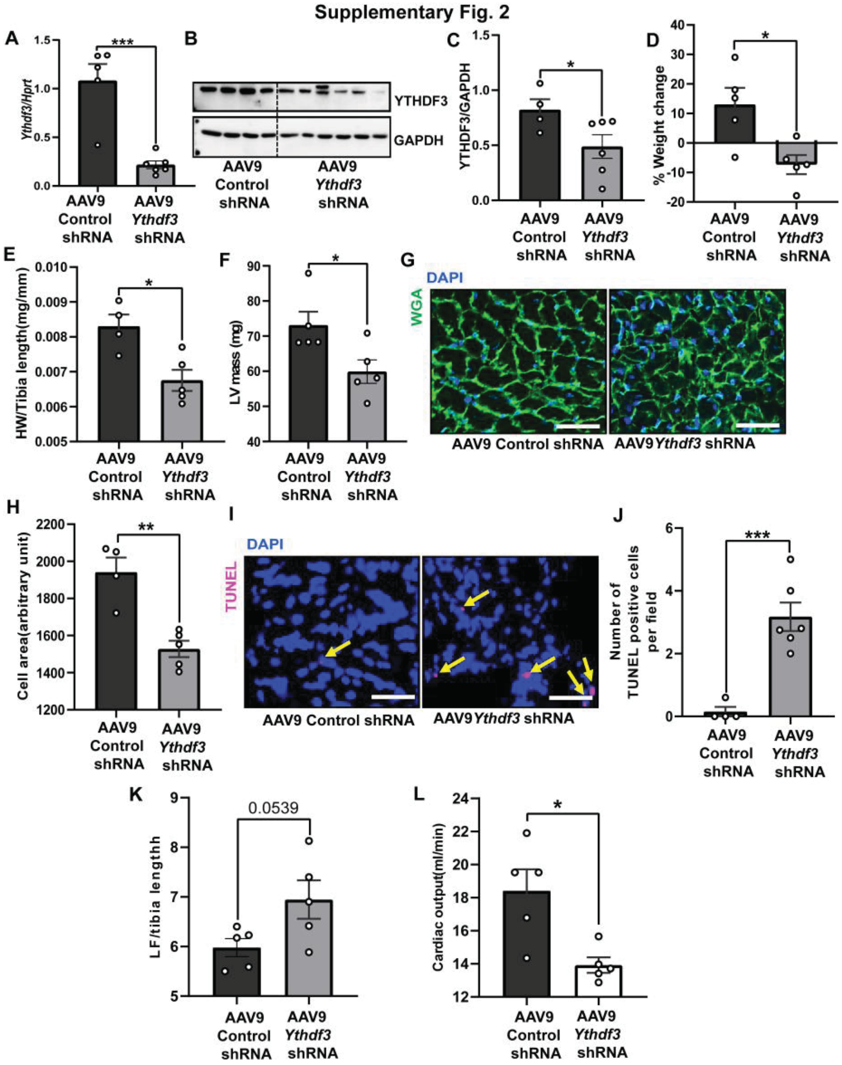


**Supplementary Fig. 2 AAV9 mediated depletion of *Ythdf3* leads to cardiac cachexia in BALB/c mice.** **(A)** Expression levels of *Ythdf3* in AAV9 Control (*n*=5) and *Ythdf3* shRNA(*n*=6) injected mice, Student t-test, unpaired (two-tailed) ***$p$=0.0005. **(B-C)** Western blot showing cardiac protein level of *Ythdf3* in animals injected AAV9-Control-shRNA and AAV9-*Ythdf3*-shRNA, Control (n=4), *Ythdf3* knockdown (n=6) Student t-test, unpaired (two-tailed) with Welch's correction *$p$=0.0497. **(D)** Percentage body weight change of animals injected with Control(n=5) and *Ythdf3* shRNA (n=5) injected mice Student t-test, unpaired (two-tailed) *$p$=0.0196. **(E)** Heart weight to tibia length ratio of AAV-Control and *Ythdf3* hearts, Student t-test, unpaired (two-tailed) *$p$<0.0116. **(F)** Left ventricular mass analyzed by echocardiography, Student t-test, unpaired (two-tailed) *$p$=0.0310. **(G-H)** Cross-sectional cardiomyocyte area of heart stained with WGA and DAPI, Student t-test, unpaired (two-tailed) **$p$< 0.0020. (**I-J)** TUNEL experiment in heart cryosection in control and *Ythdf3* depleted heart, Student t-test, unpaired (two-tailed) ***$p$=0.0008. (**K)** Lung fluid content to tibia length ratio, Student t-test, unpaired (two-tailed) $p$=0.0539 **(L)** Cardiac output measured by echocardiography Student t-test, unpaired (two-tailed) *$p$=0.0118. The scale bar represents 50μm.

#### Supplementary Fig. 3

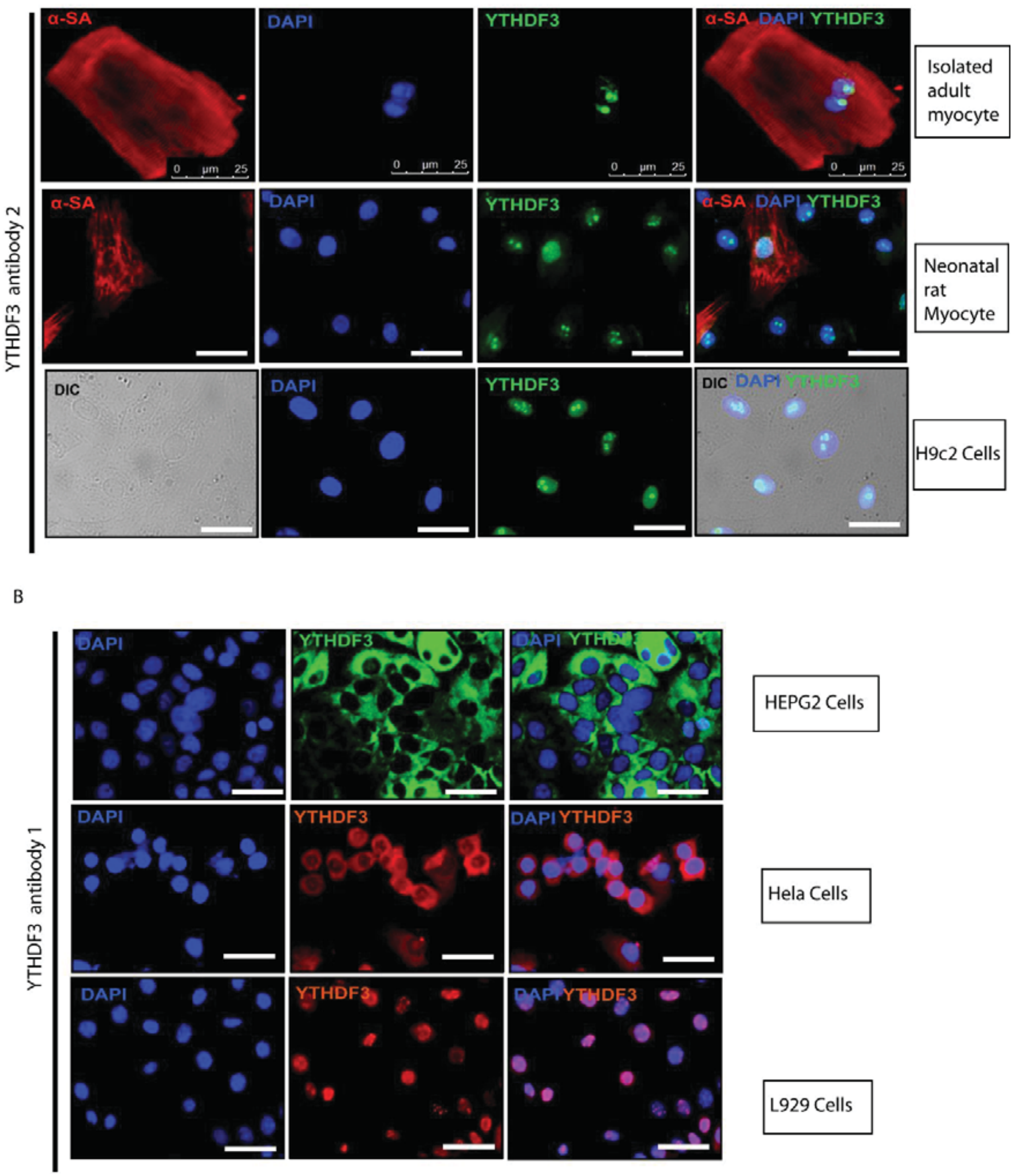


**Supplementary Fig. 3** ***Ythdf3*** **localization is cell-dependent. (A)** Localization of YTHDF3 with Abclonal antibody (2) in isolated adult mouse cardiomyocyte, neonatal rat cardiomyocyte and H9c2 cells **(B)** Localization of YTHDF3 in Human HepG2 cells, Hela cells and L929 cells with antibody1 (Invitrogen antibody).

#### Supplementary Fig. 4

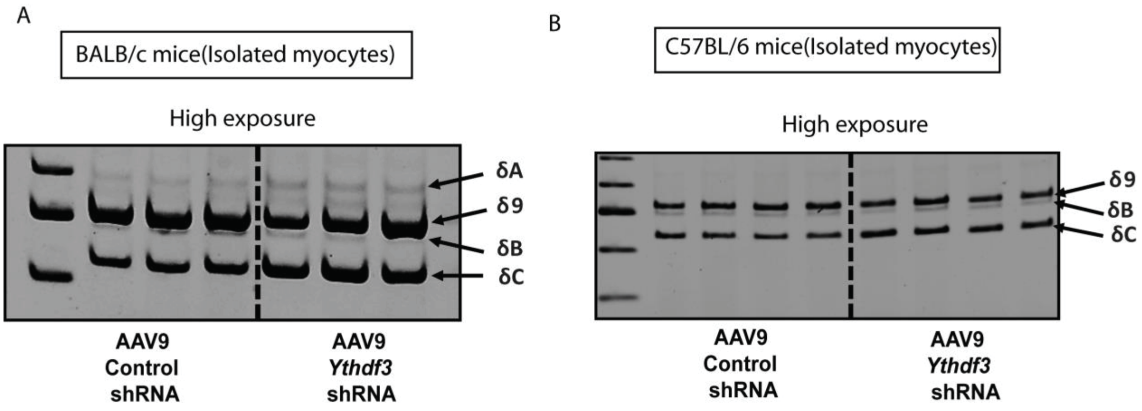


**Supplementary Fig. 4** CaMKIIδB is expressed in the heart at low levels. (**A-B**) DNA-PAGE of CaMKIIδ exons 13–18 in isolated adult cardiomyocytes from Control vs *Ythdf3* shRNA of BALB/c (**A**) and C57BL/6 (**B**) mice hearts.

### Supplementary Table 1 List of shRNAs

| **Oligos** | **Sequence 5'-3'** |
|---|---|
| Scramble shRNA FWD OLIGO | CCGGTCGTTAATCGCGTATAATACGCTCGAGCGTATACGCGATTAACGTTTTTG |
| Scramble shRNA FWD OLIGO | AATTCAAAAACGTTAATCGCGTATAATACGCTCGAGCGTATTATACGCGATTAACGA |
| Ythdf3 shRNA 1 FWD OLIGO | CCGGTACCAATGTCAGATCCATATATCTCGAGATATATGGATCTGACATTGGTTTTTG |
| Ythdf3 shRNA 1 REV OLIGO | AATTCAAAAAAACCAATGTCAGATCCATATATCTCGAGATATATGGATCTGACATTGGA |
| Ythdf3 shRNA 2 FWD OLIGO | CCGGTGGACGTGTGTTTATAATTACTCGAGTAATTATAAACACACGTCCTTTTTG |
| Ythdf3 shRNA 2 REV OLIGO | AATTCAAAAAGGACGTGTGTTTATAATTACTCGAGTAATTATAAACACACGTCCA |
| Ythdf3 shRNA 3 FWD OLIGO | CCGGTGACTAGCATTGCAACCAATCTCGAGATTGGTTGCAATGCTAGTCTTTTTG |
| Ythdf3 shRNA 3 REV OLIGO | AATTCAAAAAGACTAGCATTGCAACCAATCTCGAGATTGGTTGCAATGCTAGTCA |

### Supplementary Table 2 -List of primers

| **Primers** | **5'-3’** |
|---|---|
| *Ythdf3*_mus F | GCCACTAGCGTGGATCAGAG |
| *Ythdf3*_mus R | GCTGTTACTCTGATTTGTCTGGC |
| *Ythdf3*_rat F | TAGCCAGACAAATCAGAATAACAGC |
| *Ythdf3*_rat R | GACCAAGAAATGGAGGGGTATTTC |
| *Hprt*_mus F | GCGTCGTGATTAGCGATGAT |
| *Hprt*_mus R | TCCTTCATGACATCTCGAGCA |
| *Hprt*_rat F | GCTTTCCTTGGTCAAGCAGTAC |
| *Hprt*_rat R | GGGCATATCCAACAACAAACTTGT |
| *Camk2δ*_exon 12+13 F | TACGAGAAATTTTTCAGCAGCC |
| *Camk2δ*_exon 18 R | CCCCATTGTTGATAGCTTCAATC |
